# Sequence-only prediction of VHH nanobody epitopes and paratopes on HER2 with a frozen protein language model ( ESM2) and a structure-derived residue prior

**DOI:** 10.64898/2026.10.15.759522

**Authors:** Dharani Babu, Sundaramurthy Pandurangan

## Abstract

Mapping which antigen residues an antibody contacts (the epitope) and which antibody residues do the contacting (the paratope) is central to therapeutic antibody engineering, yet most accurate methods require a solved or predicted three-dimensional complex. We address a sequence-only formulation: predict the HER2 epitope and the single-domain antibody (VHH, or nanobody) paratope from the two sequences alone, with no three-dimensional structure prediction at any stage. We pair a frozen ESM2-650M protein language model, used strictly as a feature extractor, with a lightweight trainable module —per-chain encoders, a shared cross-attention block, and epitope, paratope and binder heads (12.2M trainable parameters) — trained jointly on three public sources: HER2 binding and blocking-competition assays (AVIDa-hHER2), residue-level antibody–antigen contacts (SAbDab2), and two reference-antibody crystal structures. The blocking-competition signal is de-confounded into a trastuzumab-minus-pertuzumab contrast whose sign encodes the HER2 domain and which is used only as a training target, never as a model input. Under a three-way, sequence-cluster-level split with a test set opened exactly once, the binder head reaches PR-AUC 0.999 and the paratope head 0.659, while residue-level epitope prediction on novel antigens remains near baseline (0.070) — a limitation the sealed split exposes rather than hides. Sequence alone recovers the competition-derived HER2 domain (the competitor class) for 87.7% of held-out clones, a leakage-free result. For HER2 specifically, a residue prior built from 25 mined HER2–antibody complexes and validated by leave-one-out reaches 0.26–0.29 residue PR-AUC once the domain is correct, roughly 3.6 times the language-model head alone. We release and report two submission variants per test clone — a fully sequence-only prediction and a competition-informed one —and document which data were used where.

## 1. Introduction

Antibodies recognise their targets through a small set of contacting residues. On the antigen, the contacted residues are the *epitope*; on the antibody, the residues that do the contacting are the *paratope*. Knowing both, at residue resolution, underpins affinity maturation, developability triage, epitope binning and the design of antibodies that avoid or deliberately share a competitor’s site. Single-domain antibodies derived from camelids (VHHs, or nanobodies) are of particular interest because their small size and single-chain format simplify engineering, but the same properties place much of their recognition in a long CDR3 loop that is hard to model.

The accurate route to an interface is a three-dimensional complex, from crystallography, cryo-EM or, increasingly, co-folding. Such structures are scarce for nanobody–antigen pairs, and for HER2 — a validated oncology target with two clinically important reference antibodies, trastuzumab (domain IV) and pertuzumab (domain II) — there are, at the time of writing, no deposited HER2–nanobody complexes at all. Predicting the interface from sequence alone, without solving or predicting a structure, is therefore both the hard case and the practically useful one. This is the task we take on, in the setting of the SEPIQ 2026 Challenge, Track A: given a VHH sequence and the HER2 antigen sequence, name the contacting residues on each side, for three held-out test clones (designated ONU11, ONU13 and ONU23) and, in principle, for any VHH.

Our contribution is threefold. First, a compact paired architecture over frozen protein-language-model features that predicts binding, paratope and epitope jointly and generalises well on the first two. Second, an evaluation protocol — a three-way, sequence-cluster-level split with a sealed test opened once — that is explicit about where sequence-only epitope prediction succeeds (at the coarse domain and competitor-class level) and where it does not (residue-level placement on genuinely novel antigens); the sealed split caught a collapse that a conventional split would have reported as success. Third, for the fixed HER2 antigen, a permitted residue-level prior assembled from known HER2–antibody structures and validated by leave-one-out, together with a transparent two-variant submission that separates the fully sequence-only answer from the competition-informed one.

We held to five self-imposed rules throughout, and they frame every section that follows:

1. **No structure prediction**. No AlphaFold, Boltz or any fold-prediction model. A protein language model (ESM2) is permitted strictly as a frozen feature extractor — it turns a sequence into per-residue numeric vectors and never predicts geometry.
2. **One generic pipeline, never per-clone rules**. The model is a single function applied identically to every VHH; the test clones are inputs, never special cases.
3. **Sequence-only at inference**. The forward pass reads only the two sequences; no assay or competition data enters it, so a clone with rich assay context and one with none receive identical inputs and graceful degradation is automatic.
4. **Test clones held completely aside**. The three test clones and every sequence in their similarity clusters are removed from training entirely.
5. **Transparency over leaderboard cleverness**. We report results as the model emits them and after any post-hoc adjustment, and we document what did not work.

## 2. Materials and Methods

### 2.1 Data

Three public sources supply the training signal; each plays a distinct role, and none is used for structure prediction — only for labels and sequence features (Table 1).

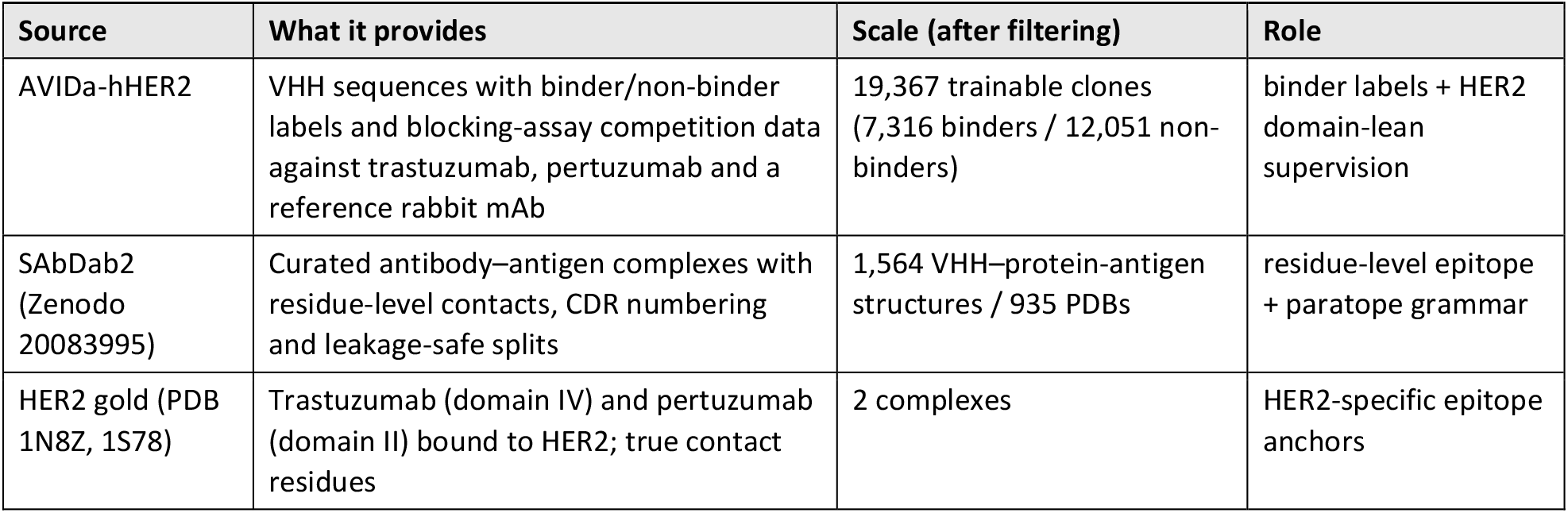

| Source | What it provides | Scale (after filtering) | Role |
| --- | --- | --- | --- |
| AVIDa-hHER2 | VHH sequences with binder/non-binder labels and blocking-assay competition data against trastuzumab, pertuzumab and a reference rabbit mAb | 19,367 trainable clones (7,316 binders / 12,051 non-binders) | binder labels + HER2 domain-lean supervision |
| SAbDab2 (Zenodo 20083995) | Curated antibody–antigen complexes with residue-level contacts, CDR numbering and leakage-safe splits | 1,564 VHH–protein–antigen structures / 935 PDBs | residue-level epitope + paratope grammar |
| HER2 gold (PDB 1N8Z, 1S78) | Trastuzumab (domain IV) and pertuzumab (domain II) bound to HER2; true contact residues | 2 complexes | HER2-specific epitope anchors |

#### AVIDa-hHER2 is the backbone

thousands of VHHs assayed against HER2, with blocking-competition experiments that reveal which reference antibody each VHH competes with — a proxy for the HER2 domain it targets. It carries binder labels and domain-level (not residue-level) epitope signal. SAbDab2 supplies the structural grammar; the legacy summary endpoint is retired, so we use the versioned machine-learning release on Zenodo, which ships pre-computed CDR numbering, antigen sequences, clusters and leakage-safe similarity splits. We derive residue-level contacts ourselves from the deposited `.cif` files with a **4.5 Å heavy-atom rule** (any non-hydrogen atom within 4.5 Å of a partner non-hydrogen atom is an interface residue), consistent with CAPRI/DockQ and the SEPIQ Voronota-LT convention; the full run processed 5,988 structures in 114 s with zero failures, at median epitope and paratope sizes of 18 and 16 residues. HER2 gold adds two crystal structures of the reference antibodies, used purely as training labels (public structures, labels only). 1N8Z pins the domain-IV footprint (trastuzumab, 19 epitope residues, 579–617); 1S78 pins the domain-II core (pertuzumab, 258–337).

#### 2.2 De-confounding the competition signal

AVIDa’s domain signal comes from blocking assays: a VHH that competes with trastuzumab likely binds near domain IV, and one that competes with pertuzumab near domain II. The catch is that trastuzumab and pertuzumab blocking are themselves correlated (Spearman ρ ≈ 0.91), because the assays share nuisance factors (expression level, avidity, plate effects); reading either raw blocking score as a domain label would mostly measure that shared nuisance. We remove the shared part with a **contrast** — the trastuzumab blocking strength minus the pertuzumab blocking strength. The nuisance common to both cancels, and what survives is the direction of the lean:

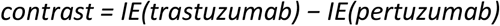

where IE (interaction energy, a log-scaled blocking strength) is taken at its most significant finite value per VHH. The sign is the domain direction — positive maps to domain IV (trastuzumab-like), negative to domain II (pertuzumab-like) — and the magnitude is the confidence: a large value is a VHH that clearly prefers one reference, near-zero is genuinely ambiguous. The de-confounded contrast agreed with an independent domain classification on 961 clones with zero mismatches (493 domain IV / 468 domain II), and crucially it is used only as a training target, never as a model input.

#### 2.3 Training manifest and label engineering

The three sources are assembled into one training manifest — one row per example, carrying the two sequences plus whatever labels and validity flags that example has. A per-example validity mask decides which loss terms fire: AVIDa rows carry binder labels (and sometimes a domain lean) but no paratope; SAbDab rows carry true residue-level epitope and paratope; HER2 gold carries epitope only. Three label-engineering decisions shape the epitope target. (i) **Flat-over-domain AVIDa targets:** AVIDa resolves epitopes only to a domain, so its epitope target is a flat distribution over the leaned domain’s residues — not a fabricated per-residue prior, which would be false precision. (ii) **Hybrid contrast-weight squash:** the |contrast| confidence becomes a per-example loss weight with a dead-zone-plus-saturation curve (weight 0 below a threshold, then a smooth ramp toward 1); direction is never squashed, only teaching strength. (iii) **Crop-around-epitope:** ESM2 has a ∼1,022-residue context window, so the few antigens longer than that are cropped to a 1,022-residue window centred on the epitope, with labels re-sliced by the same offset; HER2 (624 residues) is never cropped. The v1 manifest holds 20,767 rows (19,201 AVIDa + 1,564 SAbDab + 2 gold). The SAbDab single-domain split reproduced the dataset’s own published partition exactly (train 1,290 / held-out 274), confirming faithful assembly.

**Figure 1.**
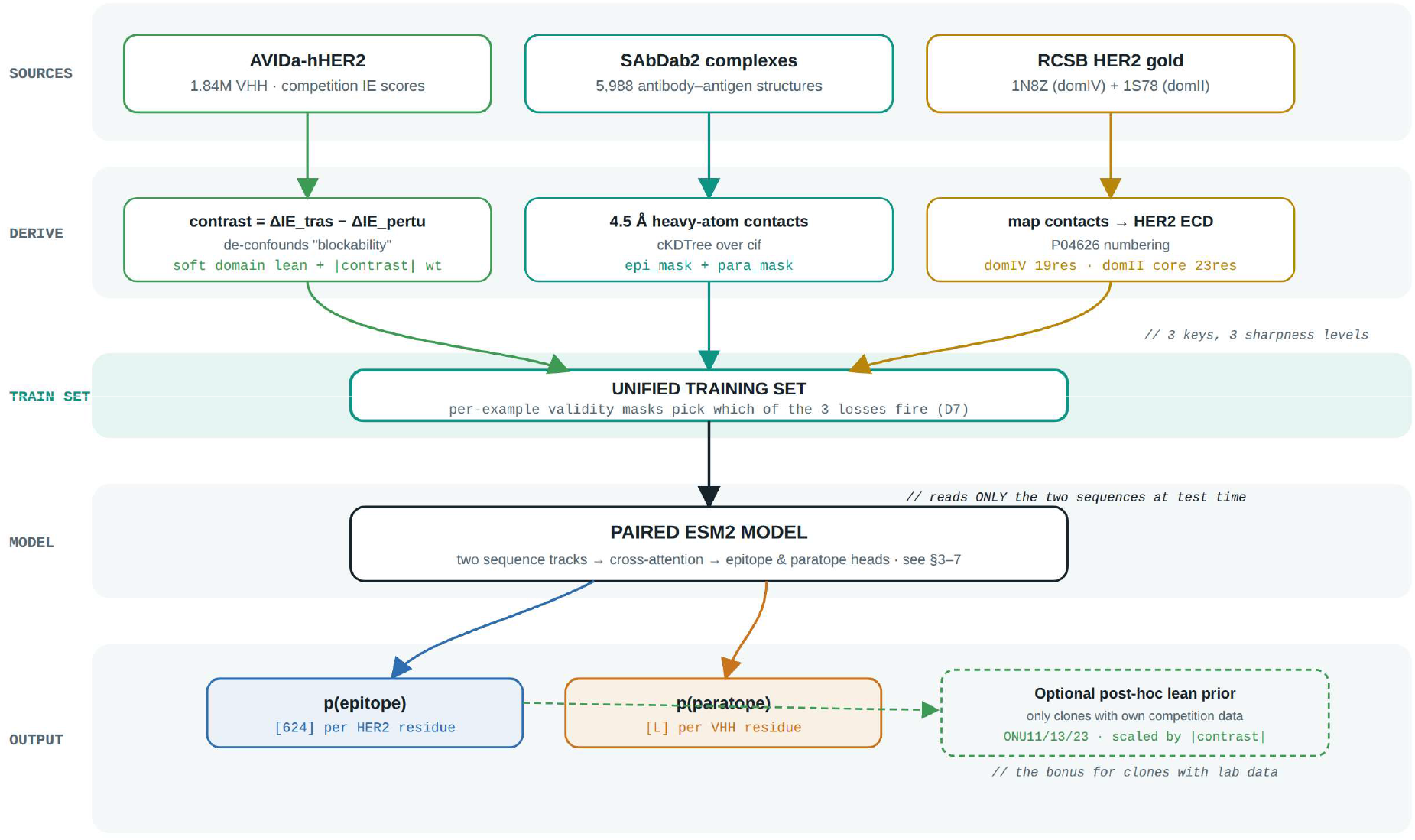
Pipeline overview. Three public sources — AVIDa competition assays, SAbDab2 contacts and the RCSB HER2 gold structures — each become a training target and converge into one unified training set governed by per-example validity masks. The set feeds the paired ESM2 model, which emits per-residue epitope and paratope probabilities; an optional post-hoc domain prior applies only to clones that carry their own competition data.

#### 2.4 Sequence features: a frozen ESM2 representation

Every sequence — antigen and VHH alike — is turned into a stack of per-residue vectors by **ESM2-650M**, run once and cached. The model is frozen: its 650M parameters are never fine-tuned, so feature extraction is a one-time cost and acts as a strong implicit regulariser (the trainable part of our model is tiny by comparison). Representations are taken from layer 33 (1,280-dimensional) and stored as fp16, with start/end tokens stripped so each cached matrix is exactly the residue grid, His-tags removed from VHHs, and keys derived from a hash of the sequence so each unique sequence is embedded once. The run embedded 20,414 unique sequences (607 antigens and 19,807 VHHs, zero overlap) into a 6.31 GB cache. CDR3 masks were derived per VHH; the v1 pipeline used a `WG-x-G` motif heuristic (92.7% coverage), and v2 adopts precise ANARCI/IMGT numbering as default (Figure 2).

**Figure 2.**
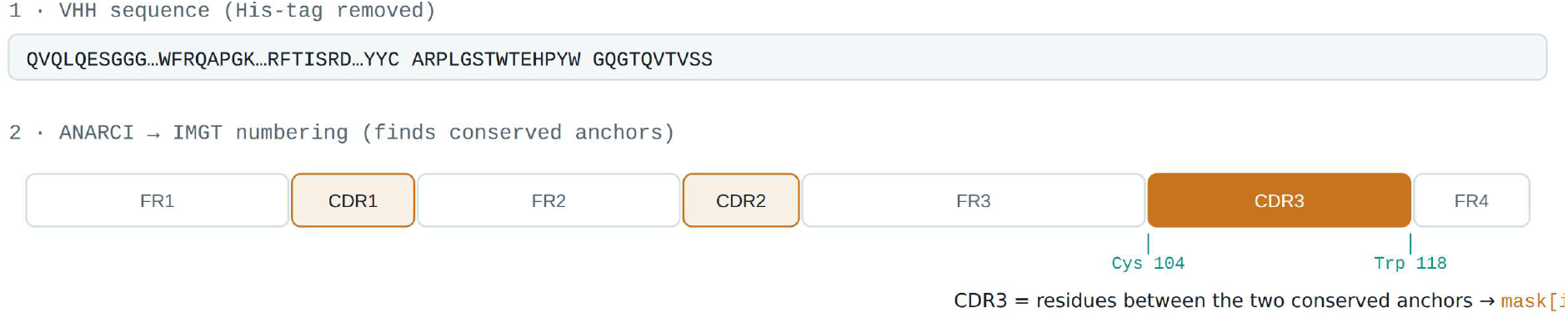
CDR3 identification. A VHH sequence is numbered by ANARCI under the IMGT scheme; framework and CDR regions are read off conserved anchors, with CDR3 taken as the segment between the conserved cysteine near position 104 and the tryptophan near 118, producing a binary CDR mask used as an input hint.

#### 2.5 Model architecture

The pipeline answers three questions at once: does this VHH bind HER2 (binder), which VHH residues bind (paratope), and which HER2 residues are contacted (epitope). Two sequences enter as frozen ESM2 features; each passes through a per-chain adapter and encoder into a shared cross-attention block, which then splits into the three heads (Figures 3 and 4). The trainable module totals 12.2M parameters (adapter 1280→384, one encoder layer per chain, two cross-attention layers, eight heads, dropout 0.1). The forward pass is sequence-only; the domain lean is never a model input — it enters training only as the epitope-head target, and an optional post-hoc prior is applied afterwards to clones that carry their own competition data.

**Figure 3.**
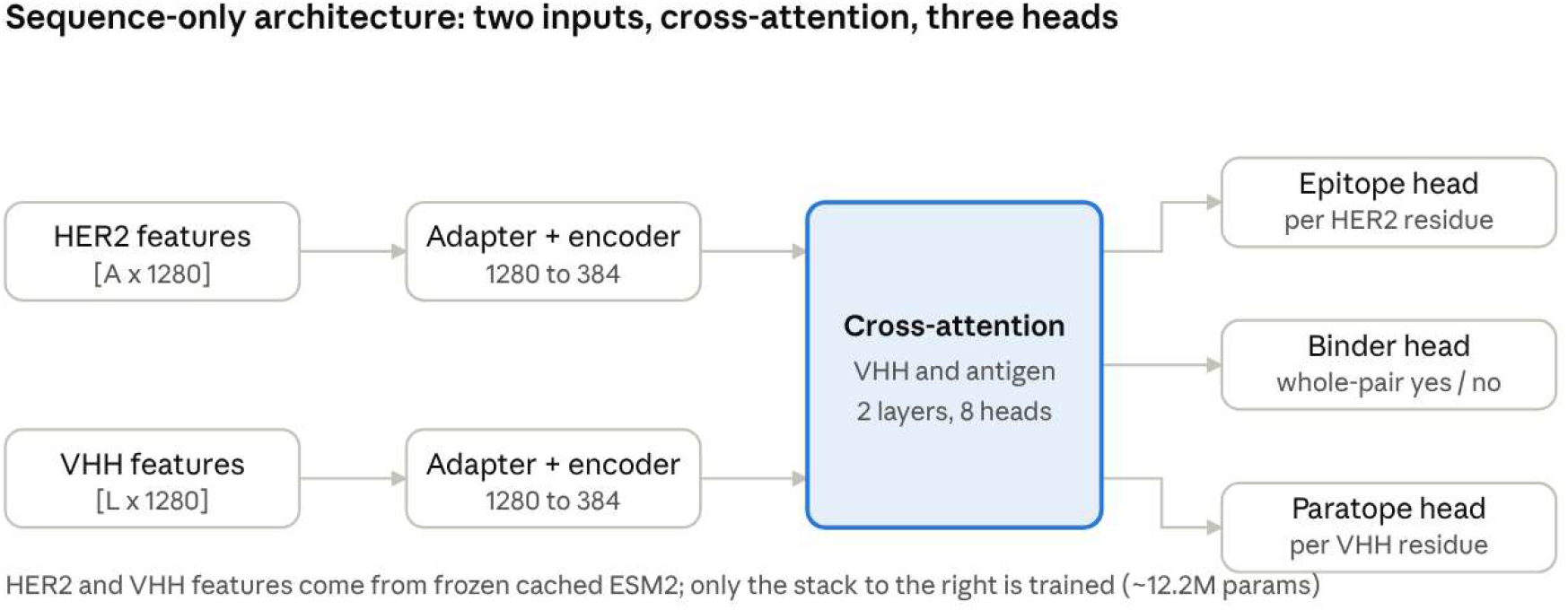
Paired model, schematically. Two sequences (HER2 and the VHH) enter as frozen features, meet in a shared cross-attention block, and split into three prediction heads (binder, epitope, paratope).

**Figure 4.**
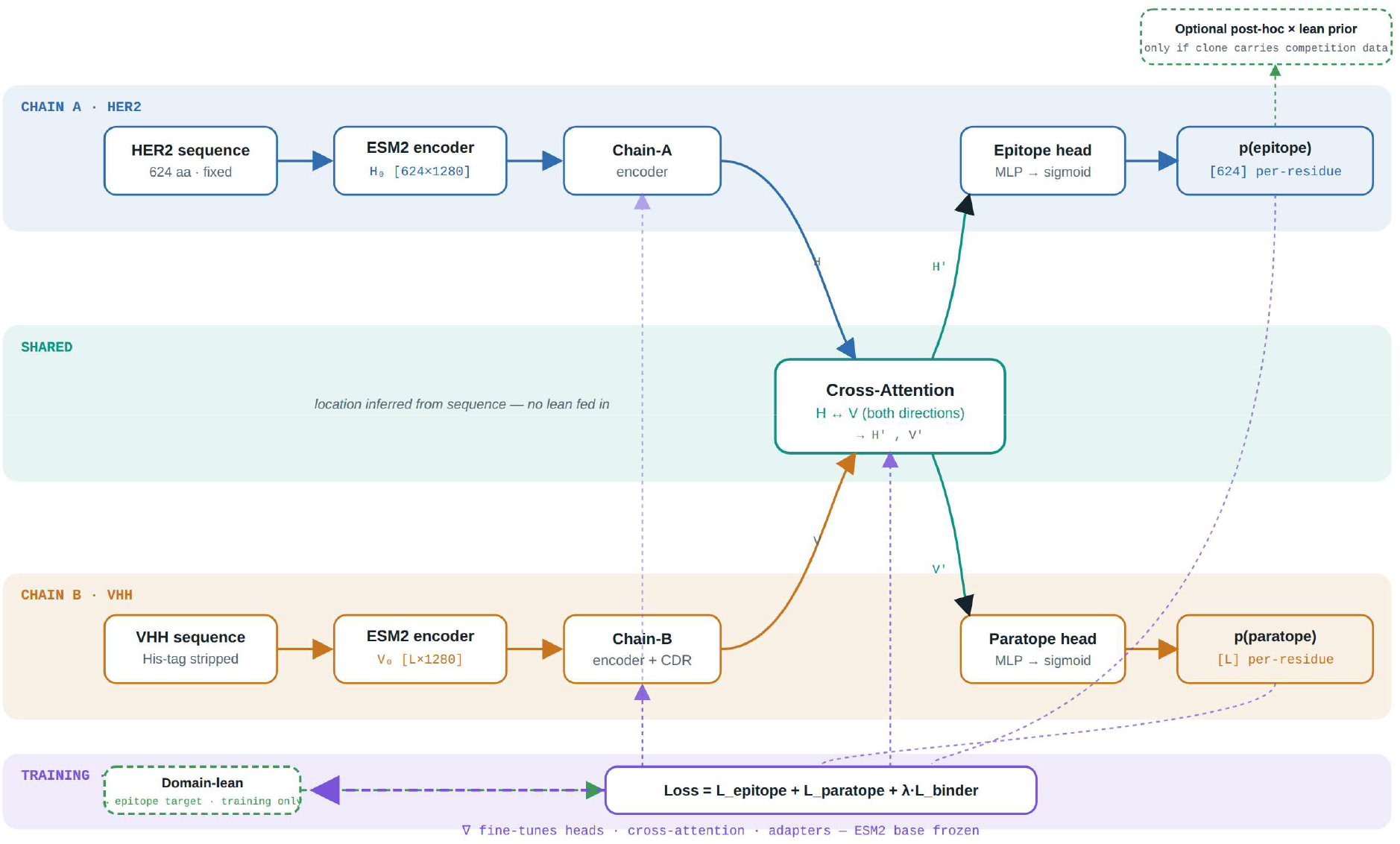
Model internals. HER2 and VHH sequences each pass through ESM2 and a per-chain encoder into the shared cross-attention block, then split into epitope and paratope heads. The forward pass is sequence-only; the domain lean feeds the epitope loss as a training target; backprop fine-tunes the heads, cross-attention and adapters while the ESM2 base stays frozen.

The cross-attention block is what makes a fixed HER2 produce a different epitope for each nanobody, from sequence alone (Figure 5). Picture a grid with HER2 residues down the side and VHH residues across the top, each cell scoring how compatible the two residues look. Reading along a row gives that HER2 residue’s epitope probability; reading down a column gives that VHH residue’s paratope probability. Swap the VHH and the bright cells move — which is why the prediction is clone-specific with no domain lean fed in.

**Figure 5.**
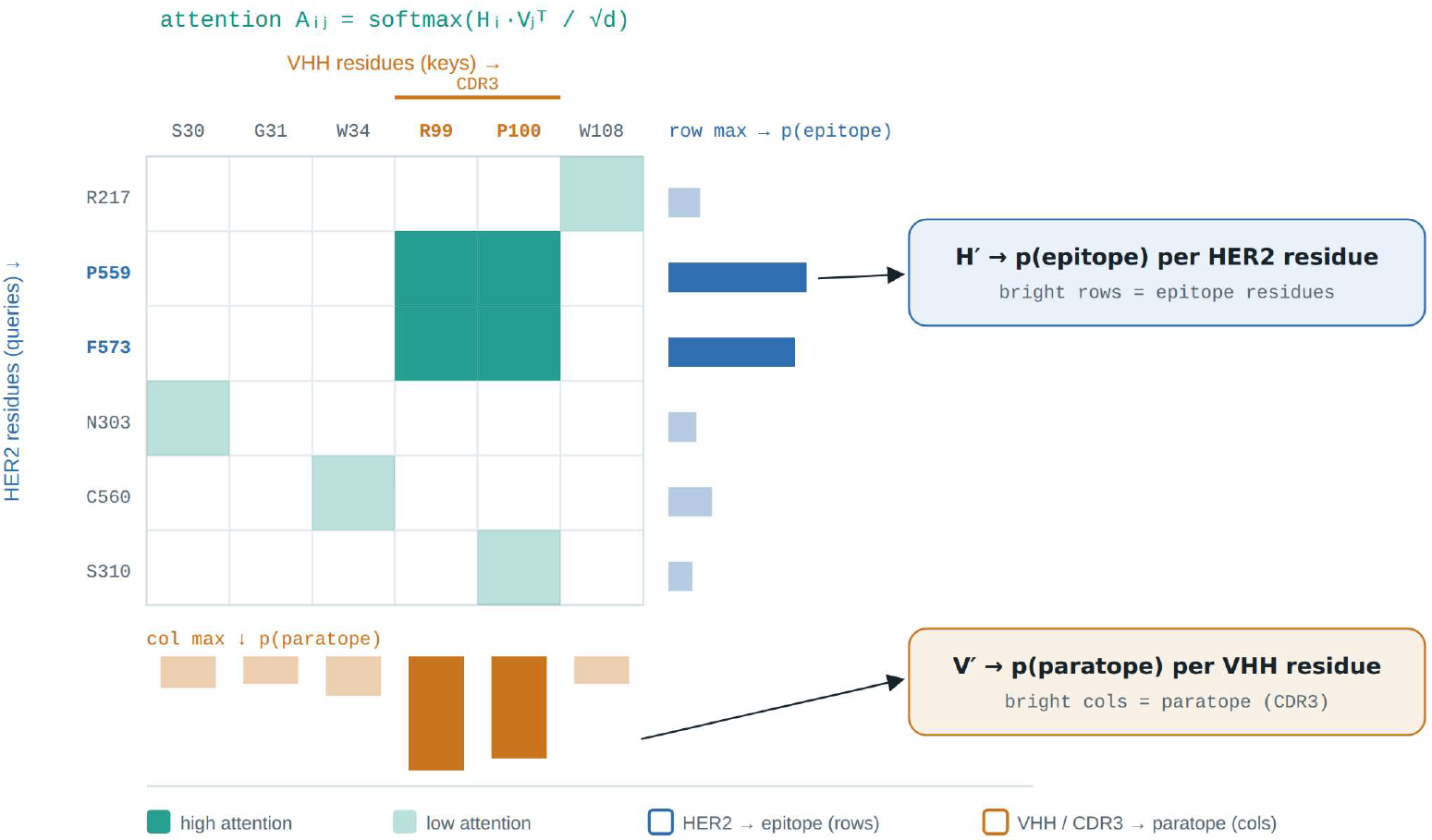
Cross-attention grid. Rows are HER2 residues, columns are VHH residues; the bright block marks a plausible contact patch where surface HER2 residues meet the CDR3 columns. Row maxima give per-residue epitope probability, column maxima give per-residue paratope probability.

#### 2.6 Training and evaluation protocol

The model is small and easy to train; the discipline is in the evaluation. We use a **three-way, cluster-level split**, every split made at the level of sequence-similarity clusters so that no cluster ever spans two splits: a training set for gradient updates, an internal validation set for early stopping and model selection, and a **sealed held-out test set opened exactly once** on the final checkpoint (v1: 18,101 / 2,012 / 654 rows). The single rule is never to tune on the set that is reported. Hyperparameters were standard and fixed (AdamW, weight decay 0.01, peak learning rate 3e-4 with warmup then cosine decay, batch size 32, gradient clipping 1.0, dropout 0.1, bf16, seed 42; binder positive-class weight ∼1.65). Because this is a multi-task model, every epoch logs train and validation loss split by head, each head’s metric against its baseline, and the generalisation gap; checkpoint selection uses a composite validation metric (binder + epitope precision + domain localisation), not raw loss.

#### 2.7 Sequence-only domain and competitor-class inference

Because the forward pass is sequence-only, the epitope prediction itself can be read as a competitor-class call without any competition data: sum the per-residue epitope probability within each HER2 domain and take the argmax — domain IV implies trastuzumab-like, domain II pertuzumab-like. This is the inverse of the post-hoc prior (which uses competition data to sharpen the epitope), and it is validatable non-circularly, because the signal being recovered — the competition-derived lean — was never a model input.

#### 2.8 A HER2 epitope atlas and exposure prior

Residue-level epitope placement on the fixed HER2 antigen is strengthened with a prior built from *known* HER2 structural data — reference data, not structure prediction. We mined the public PDB for HER2–antibody complexes (sequence identity ≥ 0.85), parsed 25 of them, and counted, per HER2 residue, how often it sits in an antibody interface (4.5 Å heavy-atom contact). That frequency is the **atlas**; a solvent-exposure mask from the free antigen’s surface area downweights buried residues. As a sanity check the atlas rediscovers the known biology unprompted — its top residues are exactly the trastuzumab (579–617) and pertuzumab (258–337) footprints. One caveat we state openly: therapeutic antibodies dominate the PDB, so the atlas encodes the common hotspots and would mislead a genuinely novel-site binder.

#### 2.9 Domain-conditioned fusion

For the test clones the chain-A (HER2) score is a domain-conditioned fusion: within the chosen domain, residues are ranked by exposure × (W_ATLAS· atlas + W_HEAD· model-head rank), and each chain is quantile-normalised to a shared scale so the pooled residue-level PR-AUC is not dominated by one chain. A sweep over W_HEAD (Section 3.7) sets W_ATLAS = 1.0 and W_HEAD = 0.5, which keeps the atlas anchoring the broad footprint while letting the model head separate same-domain clones.

## 3. Results

### 3.1 v1 sealed-test performance

On the sealed held-out test the binder and paratope heads generalised strongly and HER2 domain localisation was perfect, but residue-level epitope precision on novel antigens collapsed from a validation value of ∼0.50 to 0.055 — a failure only the sealed split could expose. Training stopped early at epoch 41 and selected the best checkpoint at epoch 31 (composite 0.830). The learning curves (Figure 6) show metrics plateauing by about epoch 13 while validation loss begins a mild upward drift around epoch 12; metric-based selection therefore keeps the model at its generalisation peak.

**Figure 6.**
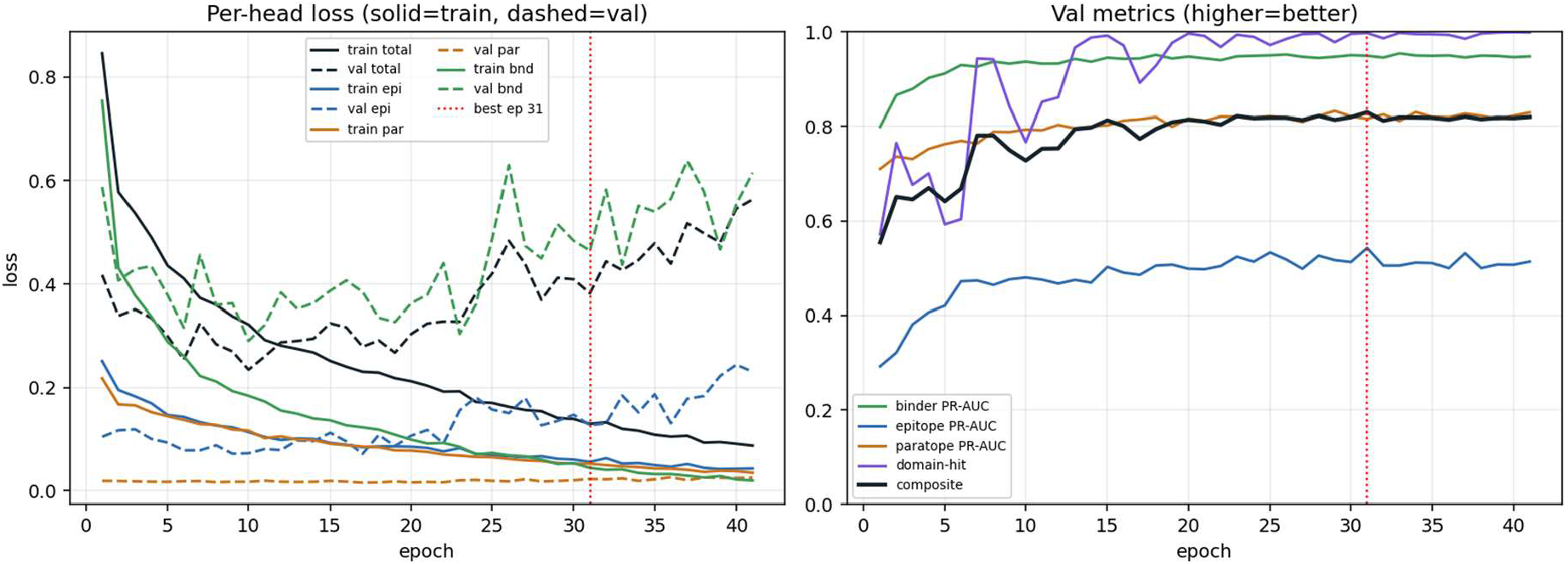
v1 training. Per-head loss (left; solid = train, dashed = validation) and validation metrics (right) across 41 epochs; best epoch 31 marked. Metrics plateau by ∼epoch 13 while validation loss drifts up from ∼epoch 12.

| Signal | Validation (peak) | Sealed test | Random baseline |
| --- | --- | --- | --- |
| Binder PR-AUC | $\sim 0.95$ | 0.998 (ROC 0.983) | 0.892 |
| Paratope PR-AUC | $\sim 0.82$ | 0.609 | 0.157 |
| Epitope residue PR-AUC | ~0.50 | 0.055 | 0.043 |
| HER2 domain-hit (median) | ~0.99 | 1.000 | — |

The diffuseness check on the 274 SAbDab test complexes (concentration ratio 0.00, precision-at-true-size 0.080, top-20 probability mass 0.851) shows the epitope head is *peaked* — 85% of its probability in 20 residues — but on unseen antigens that peak lands on the wrong residues (8% precision). It is confidently mis-localised, not smeared.

### 3.2 Antigen scarcity as the root cause

The three heads tell three different stories. Binder is a whole-sequence judgement with ∼19,000 labelled examples and transfers cleanly (ROC-AUC 0.983). Paratope — chiefly the conserved CDR3 loop — reaches ∼4× its 0.157 baseline and holds on VHHs the model never saw. HER2 domain localisation is perfect because HER2 is seen thousands of times via AVIDa, yet residue-level precision on SAbDab’s deliberately novel antigen folds is near-random: right neighbourhood, wrong house. The named cause is antigen scarcity — the SAbDab VHH set offers only ∼365 distinct antigens (∼186 joint antibody–antigen clusters), too few for the epitope head to learn transferable surface grammar, so it memorises training antigen surfaces. The architecture is sound; the epitope side is data-limited, not broken. This diagnosis triggered a pre-registered remediation (“Option B”: add antigen diversity by recovering Fab/Fv heavy chains, and add the first direct residue-level HER2 supervision).

### 3.3 v2 remediation on the same sealed set

We ran the pre-registered remediation end to end. The model, loss, optimiser, seed and sealed 654-row test set are byte-identical to v1 — only the data changed (Fv/Fab epitope rows lift antigen diversity from ∼607 to ∼2,160 distinct antigen sequences, and ANARCI/IMGT replaces the CDR heuristic). We opened the sealed box once (Table 3).

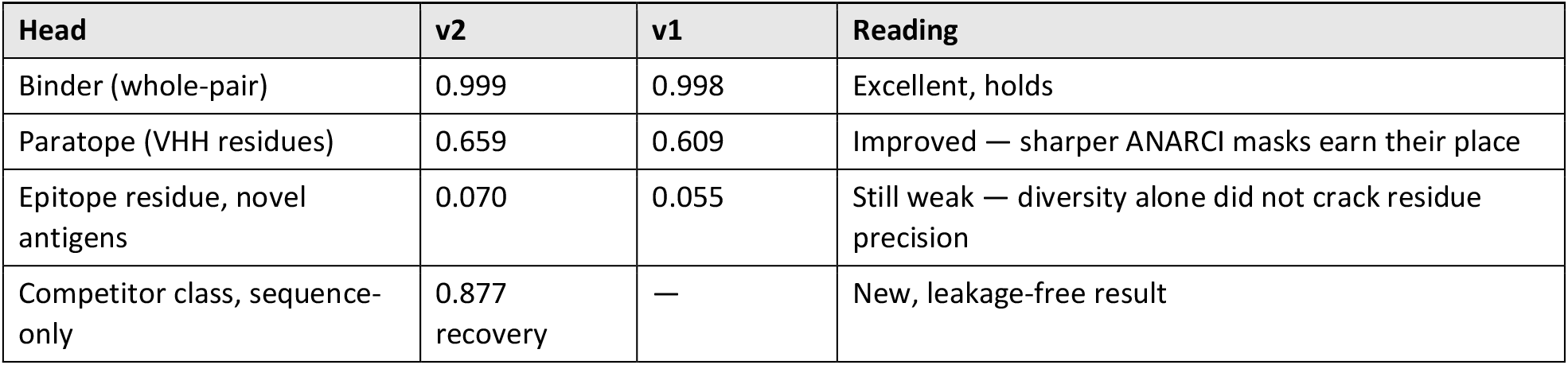

| Head | v2 | v1 | Reading |
| --- | --- | --- | --- |
| Binder (whole-pair) | 0.999 | 0.998 | Excellent, holds |
| Paratope (VHH residues) | 0.659 | 0.609 | Improved — sharper ANARCI masks earn their place |
| Epitope residue, novel antigens | 0.070 | 0.055 | Still weak — diversity alone did not crack residue precision |
| Competitor class, sequence-only | 0.877 recovery | — | New, leakage-free result |

The headline is two-sided. The diversity lever did what the architecture allows — paratope rose and binder stayed near-ceiling — but it did **not** fix residue-level epitope on novel antigens (0.055 → 0.070 against a 0.043 baseline), the hard, data-limited part of the problem, which we report plainly. The new win is at the level the task cares about: feeding only sequence, the epitope prediction recovers the competition-derived domain for **87.7%** of 57 leaned held-out clones (domain II 41/42; domain IV 9/15). Because the recovered signal was never an input, this is a non-circular test — genuine evidence that the model learned the sequence→site→competitor link. The mechanism behind both numbers is one fact: the epitope head is **domain-confident but residue-diffuse** — 99.2% of its HER2 epitope mass falls in a single correct domain, yet the top-20 residues within that domain hold only ∼16% of the mass. It knows the neighbourhood, not the street address.

### 3.4 Training dynamics: v2 versus v1

v2 re-runs the identical architecture, loss, optimiser and seed on the same sealed split, so every difference is attributable to data (Figure 7). It early-stops at epoch 29 of 39, best validation composite 0.8393 (v1: 0.8299); splits train 21,450 / internal-val 2,387 / sealed 654. Validation here is the internal-val carve, not the sealed test.

**Figure 7.**
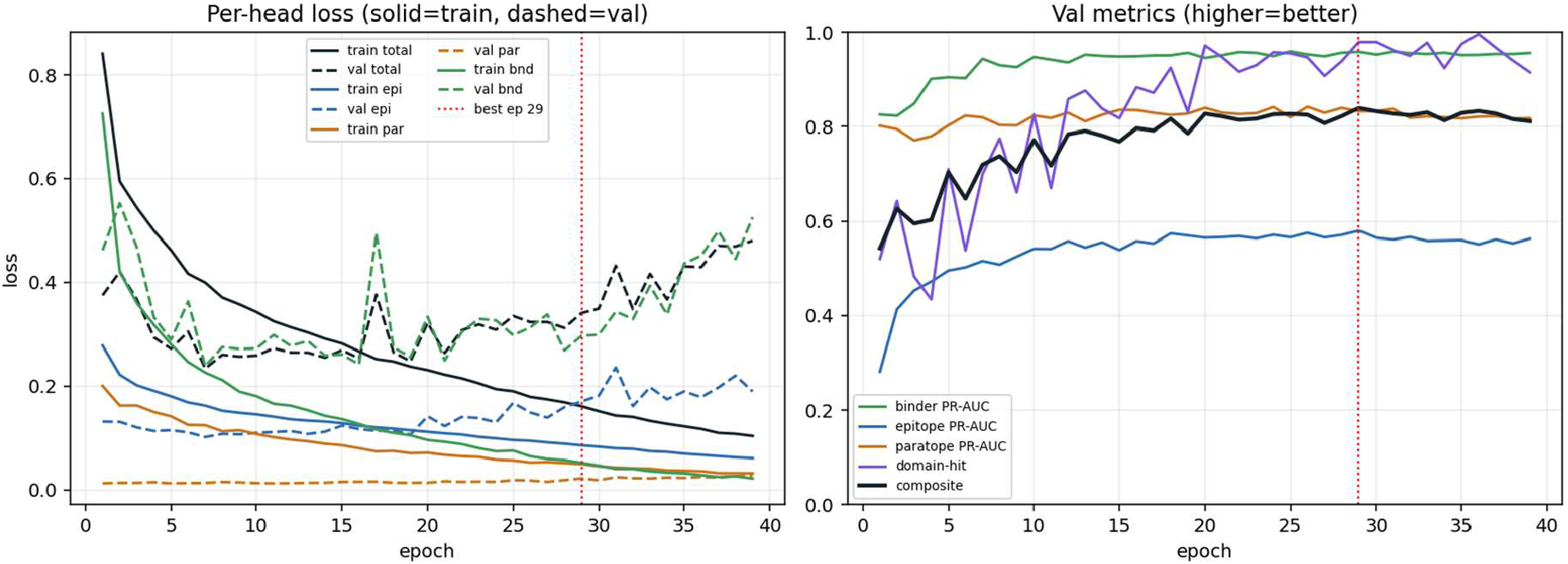
v2 training. Per-head loss (left; solid = train, dashed = validation) and validation metrics (right) across 39 epochs; best epoch 29 marked. Train epitope loss falls monotonically while validation epitope loss bottoms around epoch 7 and rises thereafter; binder, paratope and domain-hit plateau high; the composite peaks at epoch 29.

What improved:

- **Paratope PR-AUC 0.609 → 0.659** (baseline 0.157) — the clearest head-level gain, from sharper ANARCI/IMGT CDR masks replacing the `WG-x-G` heuristic.
- **Competitor-class recovery, new in v2** — the sequence-only domain classifier recovers the lean for 87.7% of held-out leaned clones (domain II 98%, domain IV 60%), leakage-free.
- **Binder PR-AUC 0.998 → 0.999** (ROC 0.991) — already near-ceiling, held cleanly; non-binders stay correctly zeroed.
- **Antigen diversity ∼607 → 2**,**160 unique sequences** — domain localisation held strong (sealed domain-share median 0.992, validation domain-hit to 0.98).

What to watch:

- **Residue-level epitope on novel antigens barely moved** (sealed 0.055 → 0.070), and residue precision-at-size actually dipped (0.080 → 0.059); diversity lifted domain localisation but not within-domain placement — which is why the HER2 submission leans on the atlas and the known-domain path, not the raw head.
- **The epitope head over-fits past the sweet spot**. Train epitope loss falls monotonically (0.278 → 0.061) while validation epitope loss bottoms at epoch 7 (0.101) and climbs to 0.19; the generalisation gap crosses zero around epochs 13–17 and reaches +0.38 by epoch 39. Composite selection takes epoch 29, before the worst — do not train longer, and do not select on raw loss.
- **The rising dashed validation loss after ∼epoch 20 is benign** binder over-confidence (validation binder loss 0.30 → 0.53 while binder PR-AUC stays ∼0.96), not a regression.
- **The right-panel validation epitope PR-AUC (∼0.58) is not the headline** — it is measured on a different internal-val carve and is not comparable across versions; the honest number is the sealed 0.070.

### 3.5 nAtlas validation by leave-one-out

There are zero HER2–nanobody complexes, so the only residue-level HER2 truth is the 25 antibody complexes themselves; we validated the atlas by leave-one-out — holding each complex out of the atlas, predicting it, and scoring it (Table 4).

| Predictor (median residue PR-AUC) | Score | vs. base |
| --- | --- | --- |
| Base rate | 0.030 | 1× |
| Model epitope head alone | 0.081 | ~2.7× |
| Atlas + exposure, model’s domain call | 0.093 | ~3× |
| Atlas + exposure, correct domain | 0.260 | ~8.7× |
| Atlas + exposure only, correct domain | 0.294 | ~10× |

Two conclusions follow. The prior is strong once the domain is right (0.294 is ∼10× the base and 3.6× the model head), and the entire bottleneck is the domain call — the jump from 0.093 to 0.260 is nothing but changing the domain source from the model to the truth. The 0.667 domain accuracy measured on these 25 complexes is a pessimistic floor, because they are conventional antibodies, out-of-distribution for a nanobody-trained model; on real VHHs the domain call is the 0.877 above. For a clone whose domain is known — because the model calls it confidently or because its competition data fixes it — the residue prior operates in the strong 0.26–0.29 regime.

### 3.6 Test-clone predictions

Each submission template is filled on the frozen checkpoint, with the atlas and exposure prior ranking residues inside the chosen HER2 domain and the paratope head ranking the VHH chain. We report two variants per clone, differing only in the HER2 domain source: a **no-competition** variant (domain from the model’s sequence-only call, fully leakage-free) and a **with-competition** variant (domain from the clone’s own competition data, an allowed input for the test clones). All three clones are confidently called binders (0.98–1.00); the domain outcomes are in Table 5.

| Clone | Model domain (conf.) | Competition lean | Agree? | Class | Footprint overlap (no-comp / with-comp) |
| --- | --- | --- | --- | --- | --- |
| ONU11 | II (0.89) | II | Yes, both weak | Pertuzumab-like | 0.85 / 0.85 |
| ONU13 | IV (0.99) | IV | Yes | Trastuzumab-like | 0.75 / 0.75 |
| ONU23 | II (1.00) | IV | No | see text | 0.00 / 0.85 |

For **ONU11 and ONU13 the two variants coincide** — the model’s sequence-only call already matches the competition data. ONU11 is the low-confidence case: its domain-II lean is a real but weak measured value (contrast −31, against +224 for ONU13 and +74 for ONU23), the clone is flagged `noise` in AVIDa, and its strongest blocker is the reference rabbit mAb; its domain-II call therefore rests mainly on the model (0.89), with competition a soft corroboration. ONU13 is the most confident prediction, a clean strong domain-IV lean that the model independently recovers. **ONU23 is where the variants diverge:** the sequence-only model is confidently domain II, but ONU23’s competition data is a clean, strong domain-IV lean — exactly the under-recovery of domain IV the sealed test flagged (9 of 15 recovered, misses landing in domain II). The allowed competition input corrects it, moving the gold-footprint overlap from 0.00 to 0.85. We therefore treat with-competition as the primary submission and report no-competition as the leakage-free companion, with the divergence shown rather than hidden (Figure 8).

**Figure 8.**
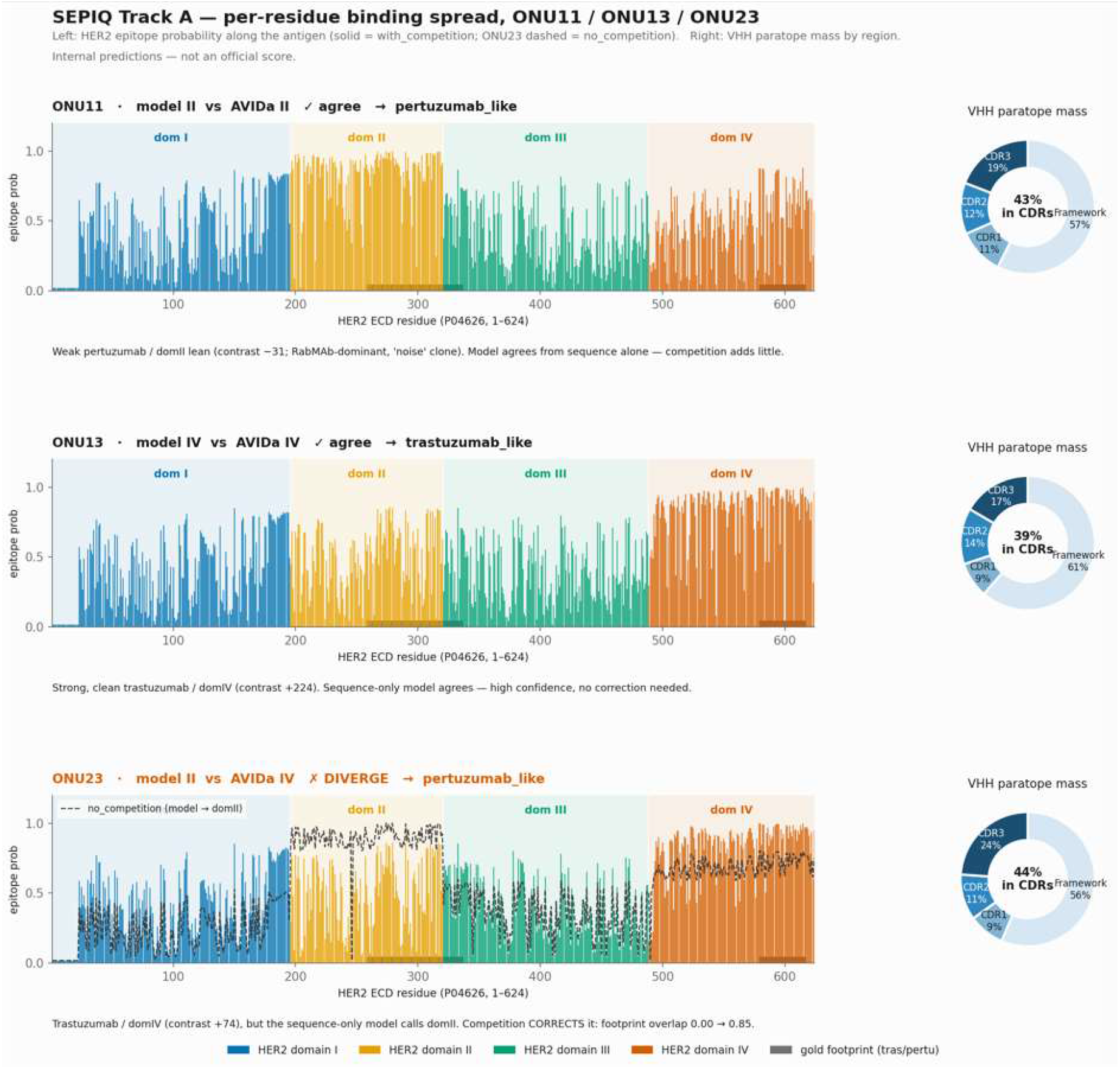
Per-clone binding maps. For each test clone, per-residue HER2 epitope probability along the antigen (domains I– IV shaded; trastuzumab 579–617 and pertuzumab 258–337 footprints marked) beside the VHH paratope’s share across CDR and framework regions. For ONU23 the model’s sequence-only call (dashed) is overlaid against the competition-corrected one.

Read across the three clones, the gain from the competition input is none for ONU11 (soft confirmation), none for ONU13 (the model was already right) and large for ONU23 (a correction) — which is the point of carrying both variants: the competition input is inert where the model is already correct and decisive exactly where it is wrong.

### 3.7 Internal evaluation and the fusion-weight choice

No HER2 residue ground truth exists for any nanobody, so the three clones are scored against proxy truth; every number here is an internal estimate, not an organiser score. The honest, held-out figure is the leave-one-out residue PR-AUC of 0.26–0.29 given the correct domain. The in-sample footprint overlaps below run higher because the atlas includes the very complexes it is scored against — a consistency check, not independent accuracy (Table 6).

| Clone | Variant | Domain used | Footprint PR-AUC (in-sample) | Top-20 mass in footprint |
| --- | --- | --- | --- | --- |
| ONU11 | no-competition | II | 0.56 | 0.85 |
| ONU11 | with-competition | II | 0.57 | 0.85 |
| ONU13 | no-competition | IV | 0.65 | 0.75 |
| ONU13 | with-competition | IV | 0.63 | 0.75 |
| ONU23 | no-competition | II (model) | 0.11 | 0.00 |
| ONU23 | with-competition | IV (competition) | 0.70 | 0.85 |

#### ONU23 is the clearest result

the no-competition variant places zero of its top-ranked mass on the true domain-IV footprint, while with-competition recovers it fully. On the fusion weight, an earlier configuration set W_HEAD = 0, which made chain A the generic atlas restricted to the chosen domain — so two clones in the same domain received identical epitope predictions, stripping per-clone signal from the antigen side. A sweep showed the model head is essentially free up to W_HEAD = 0.25 and, at 0.5, buys genuine per-clone separation (same-domain clones now differ) for only a ∼13% dip in within-domain PR-AUC measured on the out-of-distribution antibody complexes — a cost that likely overstates the true nanobody cost. The atlas still anchors the broad footprint (rank correlation ∼0.88 across the sweep); the head only re-orders the sharp top-20 end. We set **W_HEAD = 0.5**: the residue-level epitope output is clone-resolved, and the atlas-validated footprint is preserved.

## 4. Discussion

Three capabilities emerge cleanly from sequence alone: whether a VHH binds HER2, which VHH residues form the paratope, and which HER2 domain — and therefore which competitor class — the VHH leans toward. The first two generalise to unseen VHHs because they rest on abundant, transferable signal (a whole-sequence binding judgement and the structurally conserved CDR loops). The third, the 87.7% leakage-free recovery of the competition-derived domain, is the practically useful headline for an epitope-binning task, and it is honest precisely because the recovered quantity was never a model input.

The open problem is residue-level epitope placement on genuinely novel antigens, which stays near baseline (0.070). We attribute this specifically to antigen scarcity rather than to the architecture, since the paratope head — facing no such scarcity — generalises well on the identical backbone. For the fixed, heavily characterised HER2 antigen the picture is better: a structure-derived residue prior, validated out-of-sample, reaches 0.26–0.29 once the domain is correct, and the domain is exactly what the sequence-only classifier or the allowed competition data supplies. The limitation then reduces to a single missing ingredient — real HER2–nanobody structures — none of which currently exist.

Several limitations bound the claims. The HER2 atlas is dominated by therapeutic-antibody footprints and would mislead a genuinely novel-site binder, so we make no confident novel-site call. Domain IV is under-recovered by the sequence-only classifier (recalled 60%), the failure mode that makes ONU23’s unaided call wrong; where competition data exist they correct it, but a bare novel VHH leaning to domain IV may be mis-placed. The internal test-clone figures are proxy estimates, not organiser scores. The natural next step (a controlled retrain on the mined HER2 complexes with exposure-masked residue targets) is deferred until real HER2–nanobody structures or a stronger transferable signal are available, because with zero HER2–nanobody pairs such training would mostly re-learn the aggregate atlas the post-hoc prior already encodes, more transparently.

Finally, the design is deliberately generic: the same forward pass runs on any VHH with no per-clone rules, the competition-informed path simply skipped for a bare VHH. The two-variant submission — a fully sequence-only prediction alongside a competition-informed one — keeps the method’s honest unaided performance legible next to its best allowed answer.

## 5. Data and code availability

All data sources are public, versioned and citable: the AVIDa-hHER2 binding/competition dataset (challenge data can be obtained by registering for SEPIQ 2026 Challenge https://sepiq.uk/. The authors have NO right to distribute the dataset), the SAbDab2 machine-learning release (Zenodo record 20083995), and the RCSB structures PDB 1N8Z (trastuzumab–HER2) and 1S78 (pertuzumab–HER2), together with the 25 HER2–antibody complexes behind the epitope atlas (mined from the public PDB at sequence identity ≥ 0.85). Every result is regenerable from a numbered pipeline driven by a single configuration file that holds all constants (contact cutoff, crop length, split fraction, squash thresholds, learning rate, loss weights, seed 42); the data split is deterministic and a checksum manifest records the exact files used. Code is available from the author on request.

## 7. Author contributions

Dharani Babu / Sundaramurthy Pandurangan conceived the study, designed and implemented the pipeline and evaluation, and wrote the manuscript.

*References are provided in a standard numbered style; please verify the two dataset/challenge entries marked [to confirm] against their final citable form before submission.*

## Supplementary information

### S1 Pre-registered validation checkpoints

| ID | What it checks | Action if it fails |
| --- | --- | --- |
| VC1 | Epitope diffuseness: concentration ratio > 3, or precision-at-size < 0.30, or top-20 mass < 0.40, or domain-hit < 0.50 | Run the Option-B remediation (fired in v1) |
| VC2 | Whether a solvent-exposure mask sharpens the AVIDa epitope target | Use exposure-masked targets |
| VC3 | Domain-IV over-prediction | Rebalance / reweight the epitope loss |
| VC4 | Learning-rate squash sweep; binder head must not collapse | Adjust weighting (held at 0.998/0.999, no collapse) |
| VC5 | CDR3 motif heuristic vs. ANARCI numbering | Switch to ANARCI (adopted in v2) |
| VC6 | Sequence-only domain classifier recovers the competition lean | Validates the competitor-class claim (0.877) |

### S2 Key training epochs (v1)

| Epoch | Composite | Epitope PR-AUC (val) | Val loss | Note |
| --- | --- | --- | --- | --- |
| 1 | 0.555 | 0.292 | 0.417 | start |
| 13 | 0.795 | 0.475 | 0.290 | metrics plateau begins |
| 20 | 0.814 | 0.498 | 0.304 | overfit gap widening |
| 31 | 0.830 | 0.543 | 0.382 | best checkpoint (selected) |
| 41 | 0.820 | 0.514 | 0.564 | early stop |

### S3 Glossary

VHH / nanobody: a single-domain antibody fragment (∼120 residues, heavy chain only).
Epitope: the antigen (HER2) residues a binder contacts.
Paratope: the binder residues that do the contacting.
PR-AUC: area under the precision–recall curve; the right summary for rare-positive problems, read against a baseline equal to the positive rate.
Contrast: the de-confounded trastuzumab-minus-pertuzumab blocking signal; its sign is the HER2 domain direction, its magnitude the confidence.
Domain-hit: fraction of predicted epitope probability falling in the correct HER2 domain.

